# The KSHV ORF20 Protein Interacts With The Viral Processivity Factor ORF59 And Promote Viral Reactivation

**DOI:** 10.1101/2020.10.06.328641

**Authors:** D. Hoffman, W. Rodriguez, D. Macveigh-Fierro, J. Miles, M. Muller

**Affiliations:** Microbiology department, University of Massachusetts, Amherst

## Abstract

Upon KSHV lytic reactivation, rapid and widespread amplification of viral DNA (vDNA) triggers significant nuclear reorganization. As part of this striking shift in nuclear architecture, viral replication compartments are formed as sites of lytic vDNA production along with remarkable spatial remodeling and relocalization of cellular and viral proteins. These viral replication compartments house several lytic gene products that coordinate viral gene expression, vDNA replication, and nucleocapsid assembly. The viral proteins and mechanisms that regulate this overhaul of the nuclear landscape during KSHV replication remain largely unknown. KSHV’s ORF20 is a widely conserved lytic gene among all herpesviruses suggesting it may have a fundamental contribution to the progression of herpesviral infection. Here, we utilized a promiscuous biotin ligase proximity labeling method to identify the proximal interactome of ORF20, which includes several replication-associated viral proteins, one of which is ORF59, the KSHV DNA processivity factor. Using co-immunoprecipitation and immunofluorescence assays, we confirmed the interaction between ORF20 and ORF59 and tracked the localization of both proteins to KSHV replication compartments. To further characterize the function of ORF20, we generated an ORF20-deficient KSHV and compared its replicative fitness relative to wild type virus. Virion production was significantly diminished in the ORF20-deficient virus as observed by supernatant transfer assays. Additionally, we tied this defect in viable virion formation to a reduction in viral late gene expression. Lastly, we observed an overall reduction in vDNA replication in the ORF20-deficient virus implying a key role for ORF20 in the regulation of lytic replication. Taken together, these results capture the essential role of KSHV ORF20 in progressing viral lytic infection by regulating vDNA replication alongside other crucial lytic proteins within KSHV replication compartments.

## Introduction

Human herpesvirus 8, also known as Kaposi’s Sarcoma associated herpesvirus (KSHV), is an oncogenic virus that has the incredible ability of initiating and maintaining lifelong infections in its host [1]. In immunocompromised individuals, these lifelong infections can lead to the development of several malignancies including Kaposi’s Sarcoma (KS), Primary Effusion Lymphoma (PEL), and Multicentric Castleman Disease (MCD) [2–4]. Similar to other herpesviruses, KSHV undergoes a bi-phasic lifecycle that features both latent and lytic phases [1]. During latency, KSHV can persist for decades within its infected host during which time only a few viral proteins are expressed. These latency-associated proteins in concert with host machinery coordinate viral genome maintenance and replication while evading detection by host immune surveillance. However, upon reactivation into the lytic phase, the vast majority of KSHV genes are expressed in an ordered cascade, rapidly amplifying the viral genome and triggering the assembly of new viral particles. As with other herpesviruses, KSHV lytic phase replication progresses *via* a mechanism distinct from latent viral DNA replication [5]. In particular, lytic replication initiates at a different origin of replication (ori-Lyt) on the viral genome and does not occur in synchrony with the host cell [6,7]. Instead, KSHV lytic replication requires at least eight viral proteins (ORF9, ORF6, ORF40, ORF44, ORF56, ORF50 and ORFK8) including its own virally-encoded DNA polymerase (ORF9) [1,8,9] and DNA polymerase processivity factor (ORF59) [10–12]. In particular, ORF59 has been shown to be recruited to the lytic origin of replication via its binding to ORF50 (RTA) [13] which leads to the recruitment of the viral DNA polymerase and initiation of replication [10,11,14]. Therefore, KSHV lytic DNA replication is reliant on the recruitment of ORF59 and the assembly of a replication complex comprised of multiple viral lytic factors.

To accommodate the large-scale amplification of the viral genome, lytic replication is accompanied by a drastic reorganization of the nucleus. This results in major nuclear and nucleolar remodeling including relocalization of host chromatin and displacement of several host proteins such as Nucleolin and Nucleophosmin [15]. This represents one of the hallmarks of herpesvirus replication: the formation of a “bean-shaped” viral replication compartment (RC) [16–19].

Among the ~85 viral proteins encoded by KSHV, ORF20 belongs to the herpesviral core UL24 gene family, widely conserved throughout the Alpha-, Beta-, and Gammaherpesviridae [20,21]. Very little is known about the functions of KSHV ORF20 and its orthologs across the herpesvirus family. For ORF20 orthologs encoded by HSV-1 (UL24), HCMV (UL76), and MHV68 (ORF20), each have been reported to modulate cell cycle arrest and apoptosis [22–24]. Past studies have also shown that KSHV ORF20 and its orthologs UL24 and UL76 localize to the nucleoli of transfected cells [25,26],[27] with a demonstrably complex localization pattern [28]. Furthermore, UL24 was shown to contribute to nucleolar reorganization by affecting the expression pattern of key nucleolar proteins [29,30][31][25]. ORF20 mRNA expression kinetics seem to vary depending on the host cell demonstrating late gene kinetics upon *de novo* infection of primary human umbilical vein endothelial cells (HUVEC) and upon reactivation of a latently infected body cavity based lymphoma cell line (BCBL-1) [32]. However, more recently, upon lytic reactivation in endothelial cells, KSHV ORF20 was shown to be expressed as an immediate early viral gene [28]. It was also demonstrated that through its interaction with OASL, ORF20 enhances KSHV replication, possibly by regulating ribosome composition [28].

Here, we investigated the ORF20 microenvironment by proximity labeling using BioID and identified ORF59 as an ORF20 interactor. We then tracked the localization of ORF20 upon lytic reactivation and found it localizes to KSHV replication compartments where it colocalizes with ORF59. Furthermore, we found that cells infected with an ORF20-deficient virus have a severe defect in viral DNA replication, late gene expression, and subsequent virion formation. Taken together, our results indicate that ORF20 is an important regulator of KSHV lytic cycle. By placing ORF20 at the replication compartment where it interacts with KSHV processivity factor ORF59, our work support a role for ORF20 in KSHV complex lytic DNA replication.

## Results

### KSHV ORF20 proximal proteome identifies novel ORF20 interactors

To better understand the function of ORF20 during lytic KSHV infection, we used the proximity labeling method BioID to characterize the ORF20 microenvironment. We first generated an ORF20-deficient virus (ORF20_STOP_) using the BAC16 Red recombinase system [33] by mutating ORF20 genomic sequence 16 nucleotides downstream of its translation start site (**Fig 1A**). This mutation resulted in the expression of a premature termination codon. This virus was used to establish a latently infected cell line in iSLK cells that will herein be referred to as iSLK-ORF20_STOP_. Using a newly generated ORF20 antibody, we confirmed that ORF20 expression is abrogated in these cells (**Fig 1B**). Because timing of ORF20 expression is crucial to set our experimental timeline, and due to expression being previously inconsistent, we next tested when endogenous ORF20 was made in iSLK.WT cells, a KSHV positive cell line similar to the one used to establish our ORF20-deficient viral mutant. Based on observations of ORF20 expression in iSLK.WT cells relative to the ORF20_STOP_ cells, the band corresponding to ORF20 is the upper band in the doublet detected with this antibody (**Fig 1B**). Based on this pattern, we detected ORF20 protein expression as soon as 24h post lytic reactivation, making ORF20 a early gene as was recently reported [28]. We thus used this time point for all future complementation experiments (**Fig 1C**).

**Figure 1:**
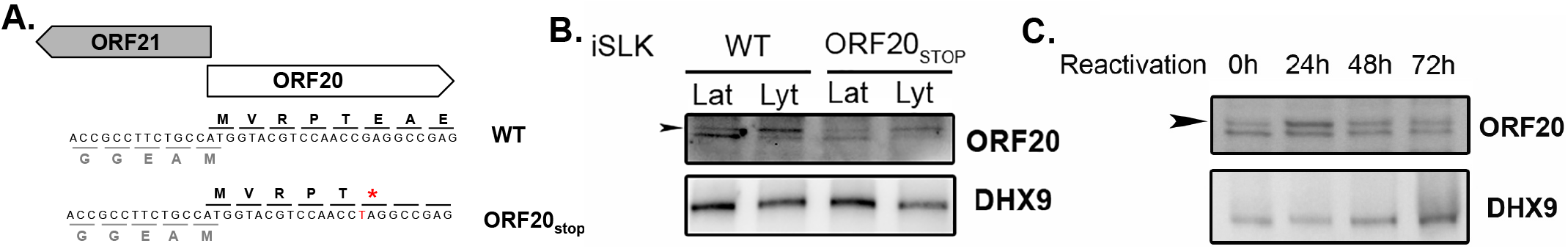
Characterization of the expression of KSHV ORF20. (**A**) Diagram showing the genomic locus of ORF20, with surrounding genes ORF21 (which partially overlaps ORF20 on the opposite strand of the KSHV genome) depicting the location of the introduced mutation. The mutation was confirmed by Sanger sequencing. (**B**) ORF20 expression was assessed by western blot in both iSLK.WT and ORF20_STOP_ cells whether in their latent phase (Lat) or lytic phase (Lyt). DHX9 served as a loading control. (**C**) Western blot showing the expression kinetics of ORF20 relative in iSLK.ORF20_STOP_ cells after reactivation with doxycycline and sodium butyrate for the indicated times. DHX9 served as a loading control.

To investigate ORF20 proximal interactome, we used the proximity labeling method BioID in which a promiscuous biotin ligase – BirA – is fused to the protein of interest. Biotinylated proteins resulting from the action of this BirA ligase can then be selectively isolated and identified by mass spectrometry [34]. We thus generated a plasmid in which the BirA_R118G_ biotin ligase was fused at the C-terminus of ORF20 which was subsequently used to rescue ORF20 expression in the iSLK-ORF20_STOP_ cells. This setup was used to identify novel ORF20 direct and indirect interactors. iSLK-ORF20_STOP_ cells were reactivated for 24h with doxycycline and sodium butyrate to trigger KSHV lytic cycle and transfected with BirA-ORF20 (or mock transfected) with excess biotin added to the media at the time of transfection. After 16h, cells were lysed and streptavidin-conjugated beads were used to isolate biotinylated proteins. The identity of the biotinylated proteins was determined by LC-MS/MS (**Fig 2A**). As shown in Supplementary Table 1, after filtering, we identified 316 high-confidence unique proteins in ORF20 microenvironment, of which 62 were previously identified with KSHV ORF20 [28]. Based on GO-term analysis on ORF20 interactors, several functional categories emerged, including several related to ribosomal regulation as previously reported (**Fig 2B**). Intriguingly, several categories placed ORF20 at the nucleosome and close to proteins regulating helicase activity. Moreover, we detected ORF59, KSHV lytic DNA processivity factor that we decided to explore further.

**Figure 2:**
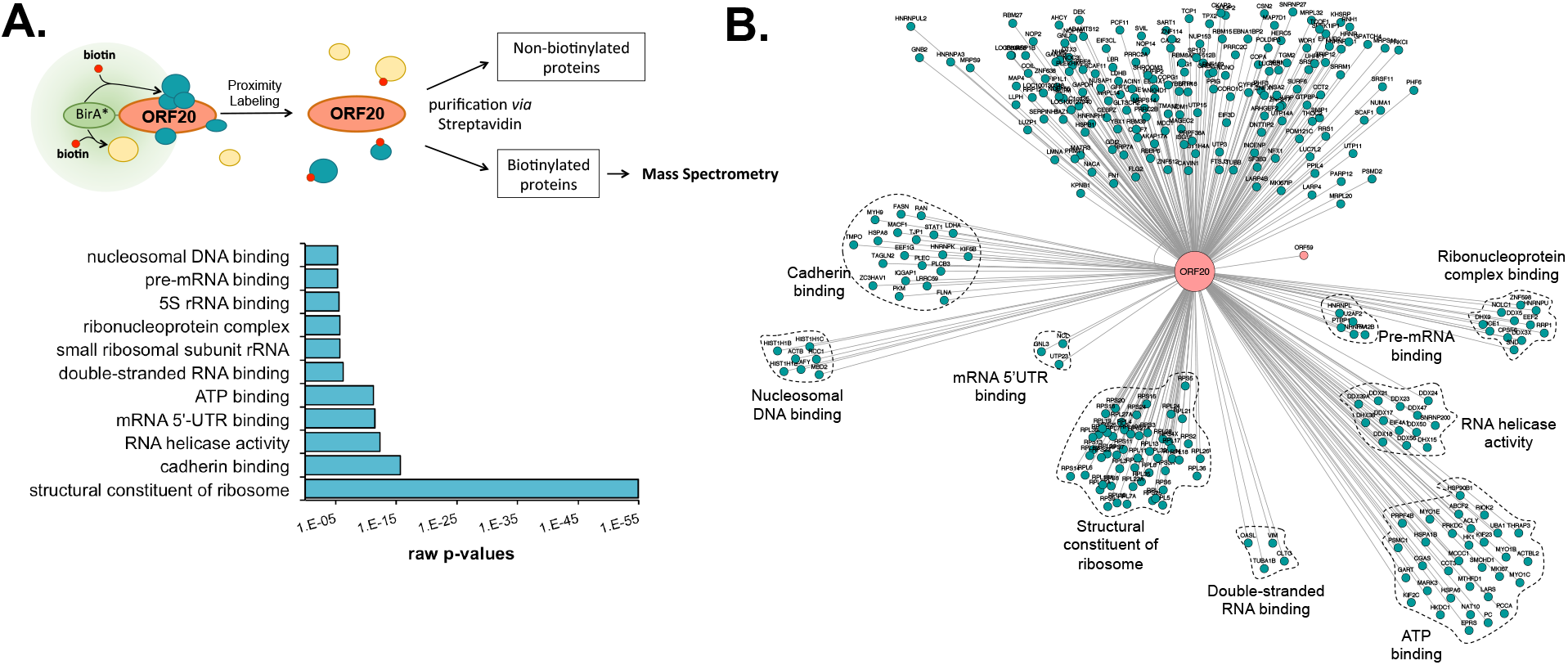
Determination of ORF20 proximal interactome. (**A**) Schematic representation of the BioID assay where ORF20 is fused to the biotin ligase BirA which will transfer a biotin to proteins coming in close proximity to ORF20. (**B**) iSLK.ORF20_STOP_ cells were transfected with ORF20-BirA and subjected to BioID and mass spectrometry. The bar graph represents the top GO-terms associated with ORF20 interacting partners and the network, generated by Cytoscape represents ORF20 proximal interactome. Proteins annotated in the categories highlighted in the bar graph are shown in clusters.

### KSHV ORF20 interacts with ORF59 and localizes in the replication compartment

To validate the interaction between ORF20 and ORF59 detected by BioID, we used co-immunoprecipitation (Co-IP): cell lysates were collected from iSLK.WT cells either in their latent or reactivated state when both ORF59 and ORF20 should be expressed. We immunoprecipitated ORF59 overnight and immunoblotted the eluates. We found that ORF59 and ORF20 interact during KSHV lytic phase (**Fig 3A**).

**Figure 3:**
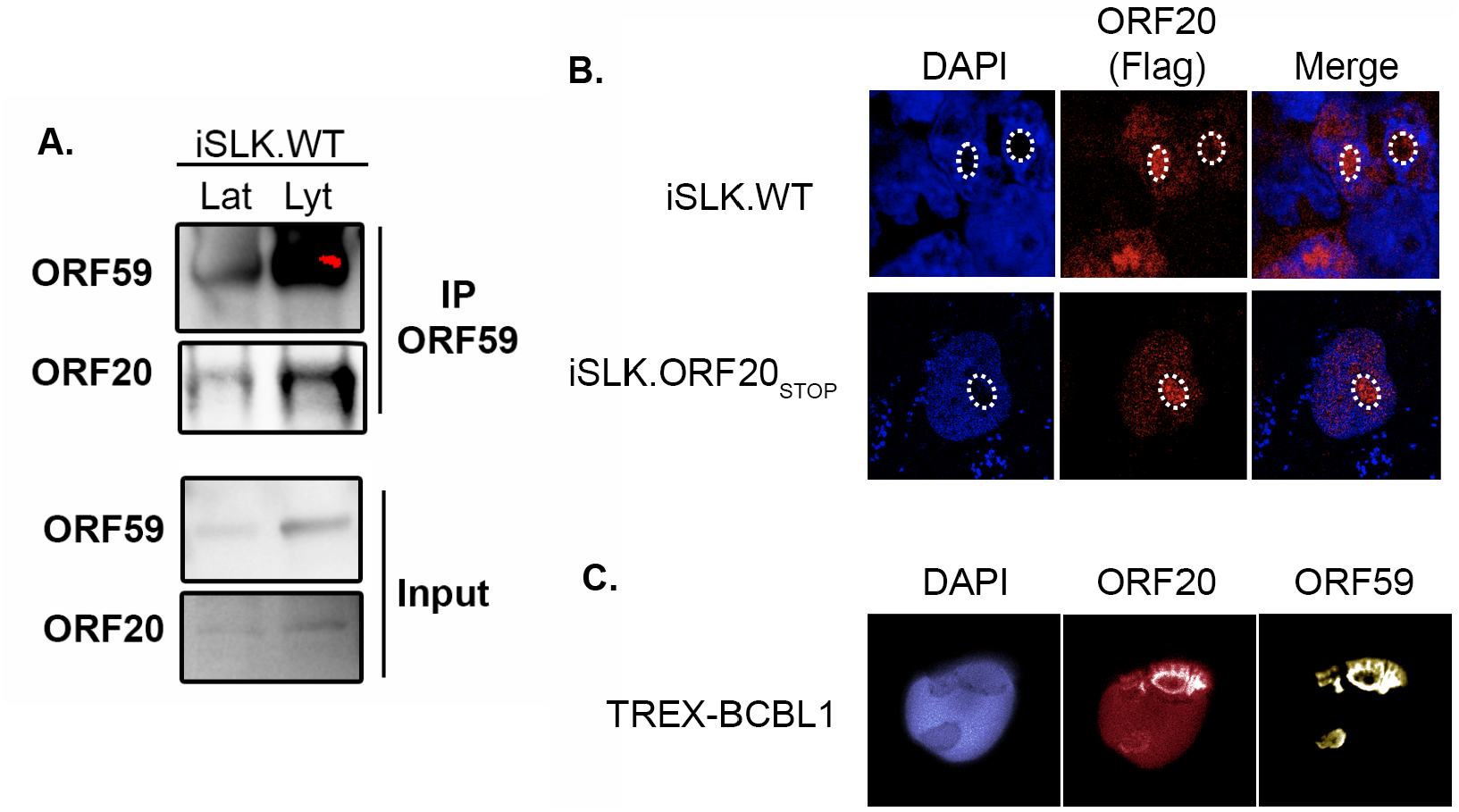
ORF20 interacts and co-localizes with ORF59 in KSHV Replication Compartments. (**A**) Lysates of latently infected (Lat) or DOX-reactivated (Lyt) KSHV-positive iSLK.WT cells were subjected to IP with an anti-ORF59 antibody, then Western blotted (WB) for ORF20 or ORF59. (**B**) Reactivated KSHV-positive iSLK.WT or iSLK.ORF20_STOP_ cells were transfected with Flag-ORF20 and 24h later were subjected to immunofluorescence assay using an anti-FLag antibody (red), and DAPI staining to identify nuclei (blue). White dotted circles denote DAPI-deficient nuclear domain representing KSHV replication compartments. (**C**) DOX-treated KSHV-positive TREX-BCBL1 cells were transfected with Flag-ORF20 and 24h later subjected to immunofluorescence assay using an anti-Flag antibody (red), anti-ORF59 antibody (yellow) and DAPI staining to identify nuclei (blue).

ORF59 has a distinctive subcellular localization during KSHV lytic cycle and localizes into replication compartments. These replication compartments are the site of the viral genome replication and are easily identifiable as dark areas in DAPI staining of host nuclei. Given that ORF20 was previously shown to localize to discreet nuclear foci, we hypothesized that ORF20 could colocalize with ORF59 at these RCs. We first investigated if we could detect a fraction of ORF20 in the RC in reactivated iSLK cells by immunofluorescence assay (IFA). In iSLK.WT cells transfected with Flag-ORF20, ORF20 expression was predominantly nuclear and distinctively present in nuclear puncta overlapping with DAPI-negative areas. Because in iSLK.WT cells, endogenous viral ORF20 expression may interfere with our Flag tagged construct, we confirmed this expression pattern in iSLK.ORF20_STOP_ cells, where we also found ORF20 expressed in DAPI negative areas of the nucleus (**Fig 3B**). Since this expression pattern is consistent with ORF59 localization patterns and our BioID and Co-IP indicate an interaction between ORF59 and ORF20, we next monitored the localization of both ORF59 and ORF20. To do so, we used the KSHV-positive TREX-BCBL-1 cells since they do not have any fluorescent marker and thus allowed us to use an additional color channel. In these cells, ORF59 and ORF20 appear to co-localize in the nucleus, in areas reminiscent of replication compartments (**Fig 3C**).

### KSHV lacking ORF20 shows defect in virion formation and late gene expression

Given that ORF20 function is poorly understood, we next wanted to better define its role during KHSV lytic cycle. Based on our data, ORF20 is expressed early in the lytic cycle and is expressed in RCs where it interacts with ORF59. Moreover, ORF20 is very well conserved in the herpesvirus family, suggesting that its function might be crucial for proper progression of viral infection so we hypothesized that ORF20 could be important for regulation of the viral replication. To address this question, we first monitored the ability of the iSLK.ORF20_STOP_ cells to produce progeny virions by supernatant transfer assay and showed that compared to iSLK.WT cells, these almost completely failed to produce any viable virions (**Figure 4**).

**Figure 4:**
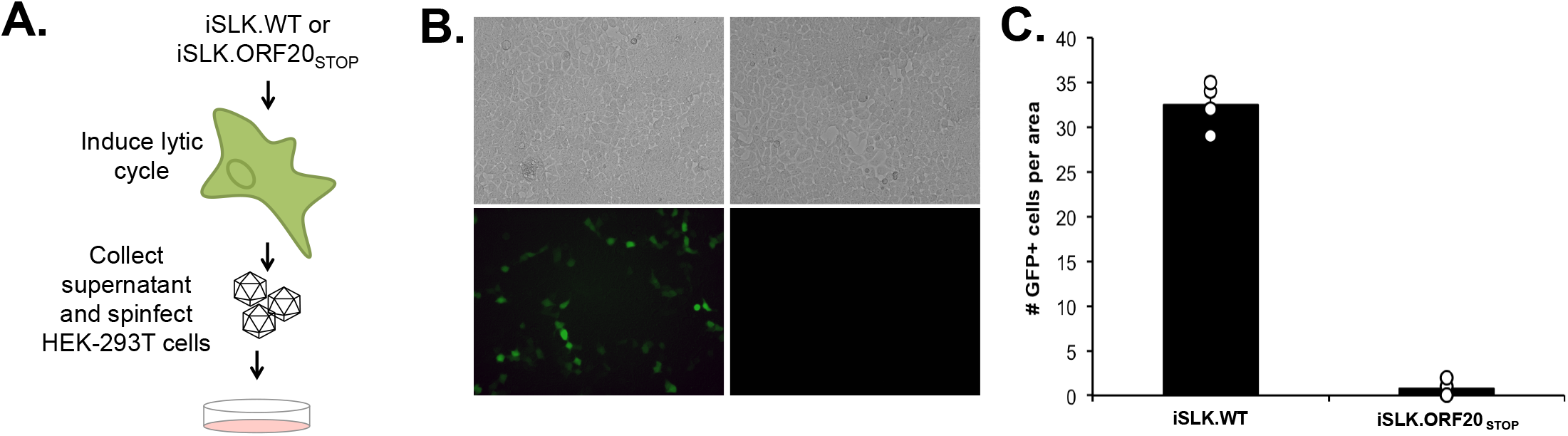
Lack of ORF20 reduces virion production. (**A**) Diagram depicting the supernatant transfer assay in iSLK cells. (**B**) Supernatant transfer assay was used as a proxy for virion production and performed as described in panel A. Infection of HEK-293T cells was monitored by contrast (top) and GFP imaging (bottom) on a fluorescence microscope. (**C**) Quantification of GFP-positive cells. Values represent four independent views of the infected cells.

Provided that the ORF20_STOP_ cells were severely deficient in virion production, we next investigated whether ORF20 expression, or lack thereof, influences viral gene expression. We investigated the expression kinetics of two viral genes for which we have antibodies: the early viral protein ORF59 and the late major capsid protein K8.1. iSLK.WT or iSLK.ORF20_STOP_ cells were reactivated and either total RNA was extracted and used for qRT-PCR or cell lysates were collected and used for immunoblot. Compared with iSLK.WT cells, iSLK.ORF20_STOP_ cells displayed a drastic reduction of K8.1 expression both at the RNA (**Fig 5A**) and protein levels (**Fig 5B**) but only a marginal effect on ORF59. Because of this differential effect on an early (ORF59) vs. late gene (K8.1) expression and because ORF20 appears to be expressed in replication compartments, we hypothesized that ORF20 could have an impact on KSHV DNA replication. Therefore, we next quantified this process in both iSLK.WT cells and iSLK.ORF20_STOP_ cells. We observed a reduction in viral DNA replication in cells lacking ORF20 expression (**Fig 5C**) suggesting that the presence of ORF20 is required for proper progression of KSHV DNA replication and consequently, late gene expression and viable virion production.

**Figure 5:**
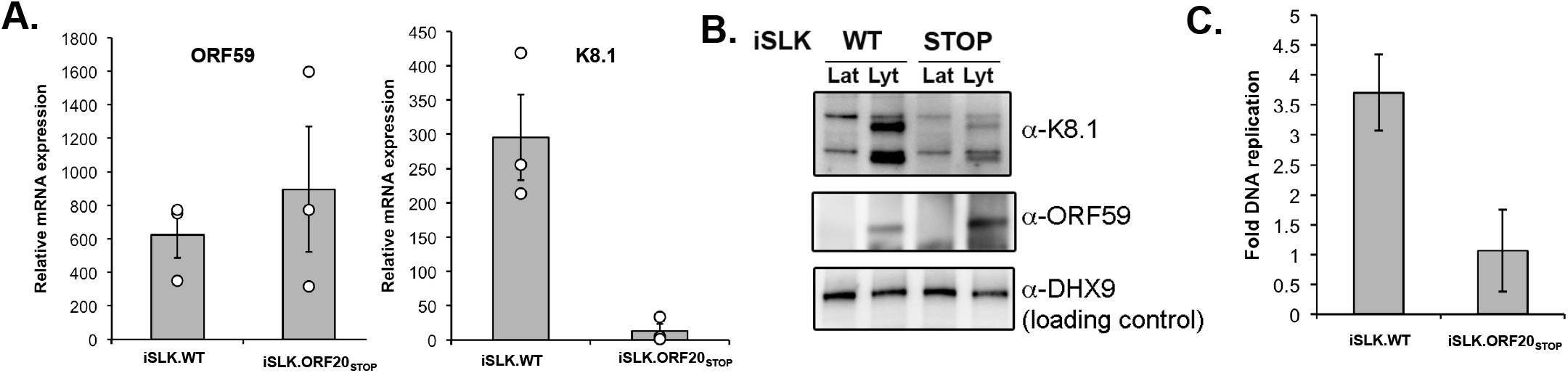
ORF20 is involved in regulating DNA replication and subsequent late gene expression. (A & B) Total RNA was extracted from unreactivated and reactivated iSLK.WT or ORF20_STOP_ cells. (A) RNA was then subjected to RT-qPCR to quantify expression of the indicated viral genes. Bars represent the relative mRNA fold change over unreactivated cells. (B) Western blot showing the expression of ORF59 and K8.1 in iSLK.WT or ORF20_STOP_ cells after reactivation with doxycycline and sodium butyrate. DHX9 served as a loading control. (C) DNA replication was measured by qPCR of the viral genome before and after reactivation of the lytic cycle.

## Discussion

In this study, we demonstrate that KSHV ORF20 localizes to replication compartments where it interacts with KSHV DNA replication processivity factor ORF59. Our approach to better define the contribution of ORF20 to KSHV lytic replication was to investigate ORF20 microenvironment by proximity labeling. Namely, we used BioID to uncover ORF20 proximal interacting partners, complementing past work done to reveal the interactome of ORF20 [28]. We confirmed many of the previously identified ORF20 protein partners, including many ribosomal factors as well as OASL as was previously reported [28]. A Go-term analysis of ORF20 microenvironment also revealed several novel putative functions for ORF20, with multiple relating to RNA binding. Since ORF20 was previously hypothesized to help bridge specific mRNA to polysomes [28], it would be interesting to further investigate this aspect of ORF20 biology. In addition, Nucleolin (NCL) was identified to be in close proximity to ORF20, which is intriguing since ORF20 ortholog UL24 in HSV-1 was shown to be directly involved in the dispersal of NCL during infection [29]. This suggests that this dispersal, which is also observed during KSHV infection [35,36], might be a conserved function of ORF20.

We also show that ORF20 has a distinctive subcellular localization that we identified as replication compartments. Of note, we also noticed that ORF20 subcellular localization was heterogeneous and we observed that ORF20 could also be more diffused in the nucleus dependent on experimental timing. This has been reported before [37] and suggests that ORF20 localization might be highly dynamic and only transiently needed at the replication compartment.

Replication compartments are enriched for many host and viral proteins involved lytic DNA replication that shuttle in and out depending on their function during this process [38,39]. Formation of replication compartments is a complex but common process for viruses: these membraneless compartments grow and coalesce over infection, using liquid-liquid phase separation and pushing host chromatin to the nuclear periphery [40–42]. Replication compartments are the site for viral genome replication, transcription, and packaging [19]. Finding ORF20 at this site thus brings up an important question: is ORF20 involved in KSHV DNA replication? Past work on ORF20 orthologs have suggested that this viral protein is important for viral replication [43], yet the precise mechanism is still unclear. However, ORF20 ortholog in MHV68 was not classified as an essential viral gene [44] suggesting that ORF20 contribution to DNA replication might vary slightly among the various herpesviruses.

One particularity of ORF20 is that it encodes a conserved PD-(D/E)XK endonuclease domain [45] [46]. This signature nuclease domain is found in a number of proteins that can cleave nucleic acids. The other known KSHV PD-(D/E)XK nuclease is the viral protein SOX, which is responsible for host shutoff during the lytic phase [47] [48] [49]. It is thus interesting to note that a second viral nuclease is encoded by KSHV and since we are placing ORF20 in the replication compartment, this nuclease activity could be important for DNA replication. The DNAse activity of SOX was previously hypothesized to be responsible for initiating DNA replication when a viral endonuclease is needed to create a nick in the origin of replication. However, this could never be proven and instead, SOX DNAse activity was shown to be important for branch resolution during replication [50]. Similarly, in HSV-1, SOX ortholog UL12 was demonstrated to be involved in recombination [51] but has not yet been shown to be the nickase necessary for rolling circle replication initiation. This therefore leaves open the question of which protein – viral or cellular – carries out this single stranded cleavage at the origin of replication during herpesviruses lytic replication. It would be interesting to test whether ORF20 PD-(D/E)XK domain is functional, and whether this viral protein contributes to this process.

Like other herpesviruses, KSHV lytic DNA replication is distinct from latent viral DNA replication both because it uses a specific lytic origin of replication and because KSHV relies of viral and not cellular factors to regulate this process. Regulation of this process is complex and extensive; more work is necessary to fully understand this crucial viral mechanism. As we understand more and more about these previously uncharacterized viral proteins, such as ORF20, we are getting closer to this goal. Given the frequently multifunctional roles of viral proteins, it will be of interest to also explore possible additional activities of ORF20, including how it may contribute to nuclear remodeling during KSHV replication and whether ORF20 has an intrinsic nuclease activity.

## Material & Methods

### Cells and transfections

HEK293T cells (ATCC) were grown in DMEM (Invitrogen) supplemented with 10% FBS. The KHSV-infected renal carcinoma human cell line iSLK.BAC16 [52] (Kind gift from Dr. Glaunsinger) bearing doxycycline-inducible RTA were grown in DMEM supplemented with 10% FBS. For reactivation, BAC16-containing iSLK cells were treated with 1 μg/ml doxycycline and 1 mM sodium butyrate for the indicated times. The KSHV-positive B cell line bearing a doxycycline-inducible version of the major lytic transactivator RTA (TREX-BCBL-1) [53] was maintained in RPMI medium (Invitrogen) supplemented with 10% fetal bovine serum (FBS; Invitrogen), 200 μM L-glutamine (Invitrogen), 100 U/ml penicillin/streptomycin (Invitrogen), and 50 μg/ml hygromycin B (Omega Scientific). Lytic reactivation was induced by treatment with 20 ng/ml 2-O-tetradecanoylphorbol-13-acetate (TPA; Sigma), 1 μg/ml doxycycline (BD Biosciences), and 500 ng/ml ionomycin (Fisher Scientific) for 48h.

The KSHV ORF20 premature Stop codon (ORF20_STOP_) mutant was engineered using the scarless Red recombination system in BAC16 GS1783 *Escherichia coli* as previously described [52], except using two gBlocks (IDT) to introduce the mutation. Each gBlock contained half of the kanamycin resistance cassette, as well as the desired mutation, and was joined by short overlap-extension PCR before being used as the linear insert in the established protocol. The BAC16 ORF20 mutant was purified using the NucleoBond BAC 100 kit (Clontech). iSLK cell lines latently infected with the KSHV ORF20_STOP_ virus were then established by coculture: HEK293T cells were transfected with 5μg of ORF20_STOP_ BAC DNA. The following day, the cells were trypsinized and mixed 1:1 with the KSHV negative iSLK-puro cells and then treated with 25nM 12-*O*-tetradecanoylphorbol-13-acetate (TPA) and 300 nM sodium butyrate for 4 days to induce lytic replication. iSLK cells were then selected using selection media containing 300 μg/ml hygromycin B, 1 μg/ml puromycin, and 250 μg/ml G418. Media were replaced every other day for ~2 weeks, gradually increasing the hygromycin B concentration until there were no HEK293T cells remaining. These cells are herein referred to as iSLK-ORF20_STOP_.

For DNA transfections, cells were plated and transfected after 24h when 70% confluent using PolyJet (SignaGen).

Supernatant transfers were carried in iSLK cells. The indicated cells were reactivated for 48h, Supernatants from induced iSLK cells were syringe filtered through a 0.45-μm-pore-size filter, and spinfected onto HEK293T cells at 1500rpm for 1.5 h at 37C. 24h later, cells were imaged on a fluorescent microscope.

### Plasmids

The KSHV ORF20 ORF was obtained from the KSHV ORFeome [54] and cloned into a pcDNA4 Nter-3xFlag vector. For the BirA-ORF20 construct, the sequence of the BirA ligase (containing a mutation -R118G- allowing it to act as a promiscuous biotin ligase) was PCR amplified from pcDNA3.1 MCS-BirA(R118G)-HA and cloned at the C-terminus of ORF20 in the pcDNA4 Nter-3xFlag vector. All cloning steps were performed using in-fusion cloning (Clonetech-Takara) and were verified by Sanger sequencing.

### RT-qPCR

Total RNA was harvested using Trizol following the manufacture’s protocol. cDNAs were synthesized from 1 μg of total RNA using AMV reverse transcriptase (Promega), and used directly for quantitative PCR (qPCR) analysis with the SYBR green qPCR kit (Bio-Rad) on a QuantStudio3 real-time PCR machine. Signals obtained by qPCR were normalized to 18S. For DNA replication calculation, iSLK cells were incubated with 5× proteinase K digestion buffer (50 mM Tris-HCl [pH 7.4], 500 mM NaCl, 5 mM EDTA, 2.5% SDS) and digested with proteinase K (80 μg/ml) overnight at 55°C. The gDNA was isolated by using a Zymo Quick gDNA miniprep kit according to the manufacturer’s instructions. DNA levels were quantified using relative standard curves with primers specific for KSHV ORF59 (5’- AATCCACAGGCATGATTGC -3’ and 5’- CACACTTCCACCTCCCCTAA-3’) or a region in the GAPDH promoter (5’-TACTAGCGGTTTTACGGGCG-3’ and 5’-TCGAACAGGAGGAGCAGAGAGCGA-3’). The relative genome numbers were normalized to GAPDH to account for loading differences and to uninduced samples to account for the starting genome copy number.

### Western Blotting

Cell lysates were prepared in lysis buffer [NaCl 150mM, Tris 50mM, NP40 0.5%, DTT 1mM and protease inhibitor tablets] and quantified by Bradford assay. Equivalent amounts of each sample were resolved by SDS-PAGE and western blotted with the following antibodies at 1:1000 in TBST (Tris-buffered saline, 0.1% Tween 20): rabbit anti-ORF20 (Yenzym) rabbit anti-DHX9/RNA Helicase A (Abcam), rabbit anti-GAPDH (Abcam). The rabbit anti-ORF59 and anti-K8.1 were used at 1:10,000 and 1:50,000 respectively and were a gift from the Glaunsinger lab. Primary antibody incubations were followed by HRP-conjugated goat anti-mouse or goat anti-rabbit secondary antibodies (Southern Biotechnology, 1:5000).

### Immunoprecipitation

Cells were lysed in low-salt lysis buffer [NaCl 150mM, NP-40 0.5%, Tris pH8 50mM, DTT 1mM, protease inhibitor cocktail] and protein concentrations were determined by Bradford assay. Equivalent quantities of each sample and at least 400ug of total protein were incubated overnight with the indicated antibody, and then with G-coupled magnetic beads (Life technologies) for 1h. Beads were washed extensively with lysis buffer. Samples were resuspended in Western blot loading buffer before resolution by SDS-PAGE.

### Immunofluorescence Assays

iSLK-ORF20_STOP_ or TREX-BCBL1 cells were grown on coverslips, and fixed in 4% formaldehyde for 20 min at room temperature. Cells were then permeabilized in 1% Triton-X-100 and 0.1% sodium citrate in PBS for 10 min, saturated in BSA for 30 min and incubated with the indicated antibodies at a 1:100 dilution. After 1h, coverslips were washed in PBS and incubated with AlexaFluor594 or AlexaFluor488 secondary antibodies at 1:1500 (Invitrogen). Coverslips were washed again in PBS and mounted in DAPI-containing Vectashield mounting medium (VectorLabs) to stain cell nuclei before visualization by confocal microscopy on a Nikon A1 Resonant Scanning Confocal microscope (A1R-SIMe). The microscopy data was gathered in the Light Microscopy Facility and Nikon Center of Excellence at the Institute for Applied Life Sciences, UMass Amherst with support from the Massachusetts Life Sciences Center.

### BioID and Mass Spectrometry

BioID was originally developed by Dr. Kyle Roux [Roux K. J., Kim D. I., Raida M., and Burke B. (2012) A promiscuous biotin ligase fusion protein identifies proximal and interacting proteins in mammalian cells. J. Cell Biol. 196, 801–810 10.1083/jcb.201112098] and makes use of a promiscuous biotin ligase: the BirA ligase. Samples were prepared as in [BioID: A Screen for Protein-Protein Interactions]. Briefly, iSLK-ORF20_STOP_ cells were seeded in 10cm plates and reactivated at the same time. 24h later, cells were transfected using PolyJet with 5ug of the ORF20-BirA construct (or mock-BirA control vector) and 50μM Biotin was added to the media. The following day, cells were harvested, lysed and affinity purification was performed using magnetic Strepavidin beads (Cell Signaling) overnight at 4C. Beads were then extensively washed and trypsin digested overnight. Samples were then cleaned up using C18 column and Mass Spectral data were obtained at the University of Massachusetts Mass Spectrometry Center using a Orbitrap Fusion mass spectrometer.

## Acknowledgments

We thank James Chambers and Steve Eyles for microscope and mass spectrometry support, respectively. We are thankful to all members of the Muller lab for helpful discussions and suggestions. We are also particularly thankful to Allison Didychuk for technical support with Bac mutagenesis.

This research was supported by the UMass Microbiology Startup funds and NIH grant R35GM138043 to M.M.

